# RADX condenses single-stranded DNA to antagonize RAD51 loading

**DOI:** 10.1101/2020.05.15.098996

**Authors:** Hongshan Zhang, Jeffrey M. Schaub, Ilya J. Finkelstein

## Abstract

RADX is a mammalian single-stranded DNA-binding protein that stabilizes telomeres and stalled replication forks. Cellular biology studies have shown that the balance between RADX and Replication Protein A (RPA) activities is critical for DNA replication integrity. RADX is also a negative regulator of RAD51-mediated homologous recombination at stalled forks. However, the mechanism of RADX acting on DNA and its interactions with RPA and RAD51 are enigmatic. Using singlemolecule imaging of the key proteins *in vitro*, we reveal that RADX condenses ssDNA filaments, even when the ssDNA is coated with RPA at physiological protein ratios. RADX compacts RPA-coated ssDNA filaments via higher-order assemblies that can capture ssDNA *in trans*. Furthermore, RADX blocks RPA displacement by RAD51 and prevents RAD51 loading on ssDNA. Our results indicate that RADX is an ssDNA condensation protein that inhibits RAD51 filament formation and may antagonize other ssDNA-binding proteins on RPA-coated ssDNA.

## Introduction

Genomic single-stranded DNA (ssDNA) is generated during DNA repair and replication. During DNA replication, for example, discontinuous synthesis of the lagging strand exposes short stretches of ssDNA that must be protected against nucleolytic degradation. Singlestranded DNA is also generated when replication forks stall at DNA lesions or as a result of cellular stress (1, 2). Stalled replication forks can generate additional ssDNA because of DNA polymerase and replicative helicase uncoupling, or due to the action of fork reversal enzymes and subsequent resection by the homologous recombination (HR) machinery (3, 4). Fork stability is maintained by diverse ssDNA-binding proteins, including Replication Protein A (RPA), the recombinase RAD51, and RADX. Together, these proteins regulate replication mechanisms to maintain genome stability at stalled replication forks (5, 6).

RPA is the major ssDNA-binding protein in eukaryotic cells. RPA consists of three heterotrimeric subunits—RPA70, RPA32, and RPA14—that collectively encode six oligonucleotide/ oligosaccharide-binding (OB)-folds to bind ssDNA with sub-nanomolar affinity (7). Among its diverse functions, RPA removes secondary structure, protecting ssDNA from reannealing and degradation, and acts as a loading platform for downstream repair proteins (8–10). One of these proteins is the recombinase RAD51. RAD51 displaces RPA from ssDNA in a cooperative binding reaction mediated by BRCA2 (8–10)(11). The ssDNA-RAD51 nucleoprotein filament then performs the homology search and strand invasion during double-strand break repair by homologous recombination (12–14). RAD51 also has multiple additional functions at replication forks, including regulation of fork reversal and protection of the reversed fork from excessive degradation mediated by exonucleases (15).

RADX was first identified via its enrichment at stalled replication forks and subsequently shown to bind ssDNA (16–18). RADX encodes three putative OB-folds with a domain organization that is reminiscent of RPA70 (Figure 1A). Biochemical studies revealed that RADX binds ssDNA via an N-terminal OB-fold cluster (16). Consistent with this observation, an OB-deficient mutant RADX does not rescue *RADXA* cells, indicating that DNA binding is essential for its cellular activities (18). Depletion of RADX aggravated the fork progression defect arising from elevated RPA expression, suggesting the balance between RADX and RPA ssDNA-binding activates is critical for DNA replication integrity (16). RADX-depleted cells exhibit excessive RAD51 activity and illegitimate recombination, suggesting that RADX is a negative regulator of RAD51 that functions at replication forks to maintain genome stability (17). A recent study also showed that RADX is involved in telomere maintenance by binding single-stranded telomeric DNA along with POT1 to antagonize RAD51 (19). The mechanistic basis of how RADX acts on ssDNA to negatively regulate RAD51 is unclear.

**Figure 1.**
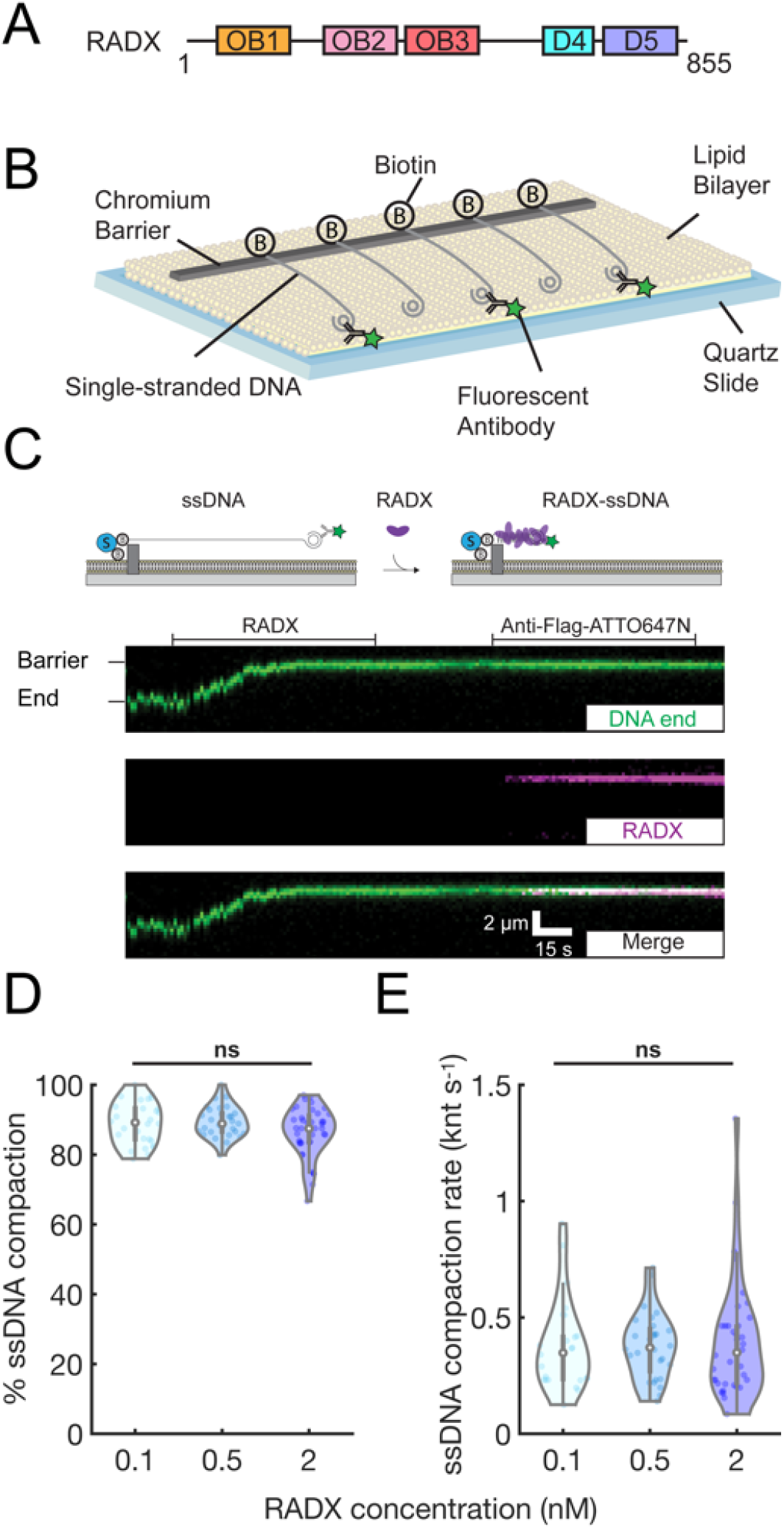
RADX condenses single-stranded DNA. **(A)** Putative RADX domain organization with 3 OB-folds. **(B)** Illustration of the singletethered ssDNA curtain assay. **(C)** Cartoon illustration (top) and a typical kymograph showing that RADX rapidly binds and compacts ssDNA. The extent and rate of compaction were monitoring via movement of the fluorescently labeled ssDNA end (green). After ssDNA binding, RADX was visualized by anti-Flag-ATTO647N (magenta). Horizontal lines indicate when RADX and anti-Flag-ATTO647N were injected. **(D)** Quantification of RADX-induced ssDNA compaction percentage. **(E)** Quantification of RADX-induced ssDNA compaction rate. Violin plots: open circles indicate the median and vertical lines show 95% quantiles of each distribution. At least 25 ssDNA molecules were measured for each condition. ns, p>0.05.

Here, we use single-molecule fluorescent imaging to dissect the functions of RADX on ssDNA substrates. RADX binds ssDNA avidly to condense both naked and RPA-coated ssDNA. Surprisingly, RADX does not displace RPA from ssDNA, but can still condense RPA-ssDNA filaments, even when RPA is present at a 100-fold excess over RADX. Furthermore, RADX inhibits RPA to RAD51 exchange on ssDNA via the formation of higher-order RPA-ssDNA structures that are refractory to RAD51 loading. We conclude that RADX preserves stalled replication forks and uncapped telomeres by antagonizing RAD51-mediated recombination via its ssDNA-condensation activity.

## Results

### RADX compacts naked and RPA-coated ssDNA

We adapted the DNA curtain assay for high-throughput single-molecule imaging of RADX-ssDNA interactions (Figure 1B). Wild type (wt) RADX encoding a single N-terminal Flag epitope was overexpressed and purified from insect cells (Supplemental Methods) (18). The ssDNA substrate was produced for single-molecule imaging via rolling circle replication of a low-complexity oligonucleotide minicircle (24). A low complexity ssDNA substrate reduces secondary ssDNA structures, which may complicate the analysis of RADX-ssDNA interactions. One end of the ssDNA was immobilized on the surface of a fluid lipid bilayer via a biotin-streptavidin linkage. The second end was fluorescently labeled with an Alexa488-labeled anti-doublestranded DNA (dsDNA) antibody that targets the 28 base pair (bp) dsDNA mini-circle (Figure 1B).

RADX rapidly compacted all ssDNA molecules to the barrier, even when injected at a concentration of 0.1 nM (Figure 1C). Labeling the RADX with a fluorescent anti-Flag antibody confirm**ed** that the protein was exclusively bound to the compacted ssDNA (Figure 1C). We measured the ssDNA compaction rate and overall degree of compaction relative to naked ssDNA by tracking the fluorescent ssDNA end. The ssDNA was 89 ± 6 % (mean ± SD) compacted at 0.1 nM RADX and the compaction rate was 0.37 ± 0.2 knt s^−1^ (Figure 1D,E). Varying RADX concentration between 0.1 nM and 2 nM did not significantly change the compaction rate or degree of compaction, suggesting that RADX binds ssDNA with sub-nanomolar affinity (Figure 1D, E). We did not observe any RADX binding to dsDNA in our experiments, as expected from prior gel-based assays (Figure S1A) (17). We also used this assay to examine RADX(OB2m), which reduces DNA binding in the strongest-affinity OB2 domain (18). RADX(OB2m) still condenses ssDNA at rates that are indistinguishable from wtRADX, indicating that strong ssDNA binding via the remaining OB-folds is sufficient for naked ssDNA compaction *in vitro* (Figures S1B, C). Taken together, these results show that RADX uses its multiple OB-folds to compact ssDNA.

Our observations with RADX are reminiscent of ssDNA compaction by *E. coli* SSB (26). SSB binds ssDNA as a homotetramer with multiple binding modes that can be experimentally defined by the NaCl concentration (24, 27, 28). SSB can compact ssDNA via wrapping of the ssDNA around the tetramer core and also because of neighboring SSB tetramer interactions (29). Therefore, we tested whether RADX-mediated ssDNA compaction is also regulated by changes in NaCl concentration. In these experiments, RADX was first pre-assembled in ssDNA in 100 mM NaCl and the imaging buffer was switched to 10 mM NaCl (Figure S2A) or 300 mM NaCl (Figure S2B). However, we did not observe any NaCl-dependent changes in ssDNA compaction; RADX-ssDNA filaments were insensitive to NaCl concentrations between 10 mM to 300 mM. However, 1 M NaCl can dissociate RADX from ssDNA and resolve the condensed complexes back to full-length ssDNA molecules (Figure S3A). These results suggest that RADX-mediated ssDNA compaction mechanisms are distinct from other well-studied SSBs.

### RADX condenses RPA-ssDNA filaments at physiological protein ratios

Cellular ssDNA is rapidly bound by RPA, the most abundant ssDNA-binding protein in human cells (about 4 million complexes per cell) (5, 30). In contrast, semi-quantitative immunoblots were used to estimate that there are ~50,000 RADX molecules per cell (15). Moreover, RADX recruitment to stalled replication forks occurs over tens of minutes and the interplay between RADX and RPA is important for fork stability *in vivo* (16). Thus, we next investigated how RADX interacts with RPA-coated ssDNA curtains.

Single-tethered ssDNA curtains were first pre-coated with human RPA-GFP and then incubated with RADX (Figure 2A). A C-terminal RPA70-GFP fusion does not disrupt RPA functions *in vitro* and in cells (22, 31). RADX still compacted the ssDNA, even when the substrate was pre-coated with saturating RPA (Figure 2A). Fluorescently labeled RADX co-localized with the RPA on condensed ssDNA filaments without significantly decreasing the RPA fluorescence intensity, indicating that RADX cannot completely displace RPA from ssDNA (see below). Low RADX concentrations only partially condensed RPA-ssDNA and at a slower rate than the corresponding naked ssDNA; 2 nM RADX was required to fully condense RPA-ssDNA (Figure 2B). The addition of 1 M NaCl removed both RADX and RPA, restoring the ssDNA substrate back to its fully extended form (Figure S3A). Surprisingly, 2 nM RADX(OB2m) could not condense RPA-ssDNA. OB2 is thus required for RADX condensation of RPA-coated ssDNA *in vitro*. This observation also explains the cellular defects observed with the RADX(OB2m) mutant (Figure S3B, C) (18). In addition, SSB-coated ssDNA is also completely compacted by RADX (Figure S3D, E), indicating that RADX does not require specific RPA interactions for this activity. We conclude that RADX likely competes with RPA and other SSBs for free ssDNA sites and that a subsaturating concentration of RADX is still sufficient to collapse ssDNA-RPA filaments.

**Figure 2.**
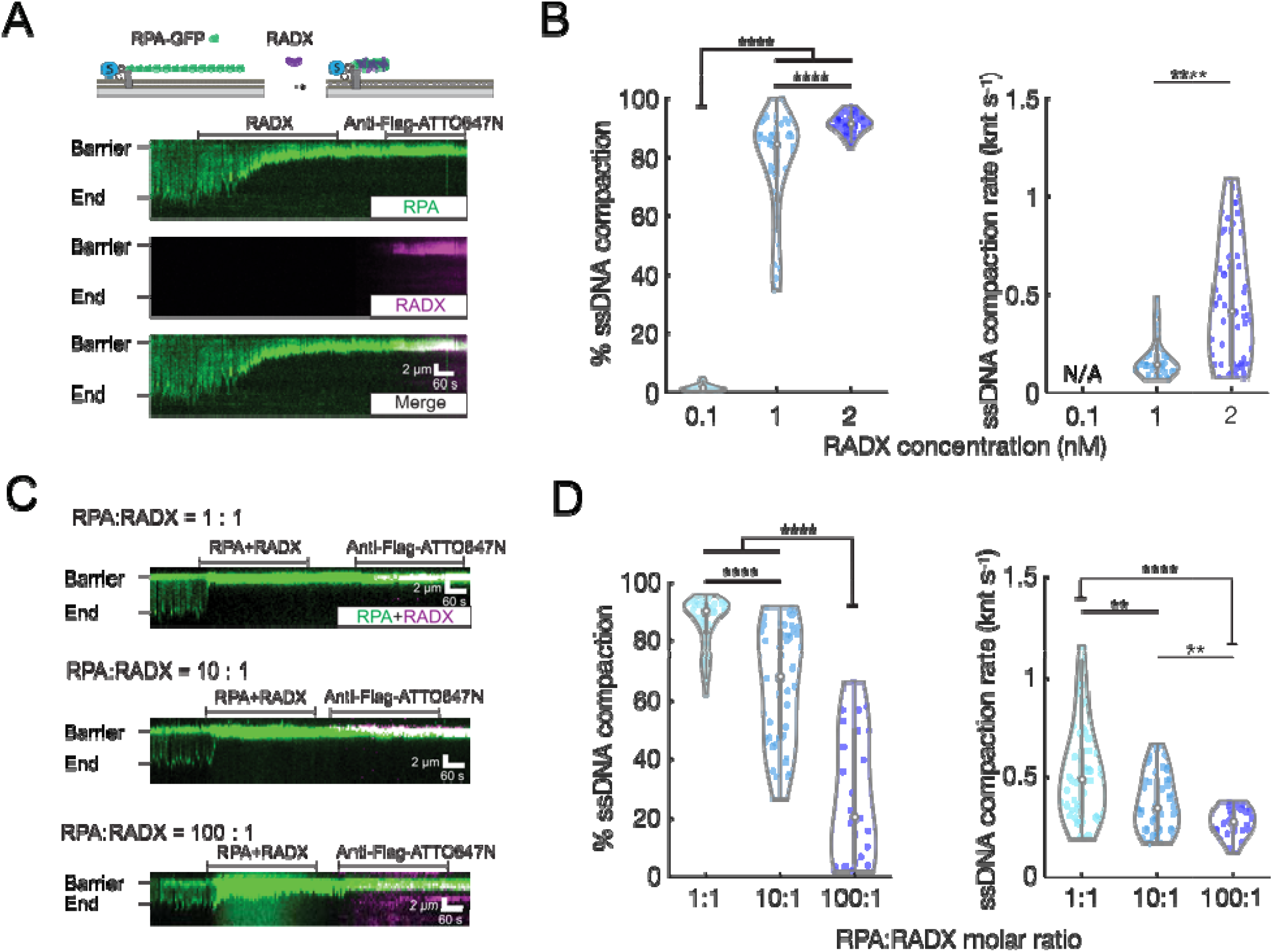
RADX condenses RPA-coated ssDNA. **(A)** Illustration (top) and a representative kymograph of RADX condensing RPA-GFP-coated ssDNA. **(B)** Quantification of RADX-induced RPA-ssDNA compaction. At least 22 ssDNA molecules were analyzed for each condition. ns, p>0.05, *, p<0.05, **, p<0.01, ***. P<0.001, ****, p<0.0001. **(C)** Kymographs of ssDNA compaction in the presence of 1:1, 10:1, and 100:1 molar ratios of RPA-GFP and RADX. **(D)** Quantification of RADX-induced ssDNA compaction at different RPA to RADX molar ratios shown in (C). At least 25 ssDNA molecules were analyzed for each condition.

We next assayed whether RADX can still condense ssDNA when it is pre-mixed with RPA at protein ratios that mimic the relative concentrations in cells (1:1, 10:1 and 100:1 RPA to RADX). In these experiments, the RADX concentration was fixed at 2 nM and the RPA concentration was increased up to 200 nM. RADX significantly compacted RPA-coated ssDNA at a 1:1 ratio (Figure 2C, top). As expected, dual-color fluorescent imaging confirmed that both RADX and RPA are present on the condensed ssDNA molecules at all RPA:RADX ratios (Figure 2C). The extent and rate of ssDNA compaction decreased with increasing RPA concentration (Figure 2D). However, RADX still condensed the ssDNA by 27 ± 20 % (mean ± SD) at the more physiological 100:1 RPA to RADX ratio. The compaction rate also decreased from 0.5 ± 0.2 knt s^−1^ to 0.3 ± 0.1 knt s^−1^ (mean ± SD) as the RPA concentration increased. In sum, sub-stoichiometric concentrations of RADX can still condense RPA-coated ssDNA filaments. RADX is ~100-fold less abundant than RPA in cells, but its recruitment to stalled replication forks and high affinity for ssDNA is sufficient to compete with RPA for ssDNA binding and to condense the nascent ssDNA gaps that occur at stalled replication forks.

### RADX bridges non-complementary DNA sequences via proteinprotein interactions

We reasoned that RADX condenses ssDNA via intramolecular association of RADX monomers into higher-order assemblies that capture ssDNA loops. To test whether RADX can self-associate intramolecularly on an extended ssDNA substrate, we prepared double-tethered RPA-coated ssDNA curtains (Figure 3A). In these assays, the ssDNA is first coated with wtRPA and then both ends of the RPA-ssDNA filament are captured between two chromium features in the microfluidic flowcell (32). The double-tethered ssDNA-RPA filament remains extended in the presence of RADX without any additional buffer flow (Figure 3A). We also purified a RADX construct that replaces the N-terminal Flag epitope with an N-terminal Maltose Binding Protein tag (MBP-RADX). Importantly, MBP-RADX and Flag-RADX both condense naked (Figure S4A, B) and RPA-coated ssDNA to the same extent and with similar rates (Figure S4C, D). We then differentially labeled the two RADX constructs with fluorescent anti-Flag or anti-MBP antibodies. Injecting 2 nM Flag-RADX (labeled with Alexa488-antibodies) into the flowcell resulted in 5 ± 2 (mean ± SD) puncta per ssDNA molecule, confirming that RADX doesn’t fully displace RPA from the ssDNA. A second injection of 2 nM MBP-RADX (labeled with a QD705-antibodies) showed that 92 ± 5 % (mean ± SD; N=113 molecules) of all Flag-RADX puncta recruited a fluorescent MBP-RADX (Figures 3B, C). Similarly, 79% of all MBP-RADX puncta co-localized with Flag-RADX. Since RPA is not replenished in these assays, MBP-RADX—which was injected tens of minutes after Flag-RADX—may encounter additional patches of naked ssDNA that produce RADX clusters. We conclude that RADX assembles into multi-protein patches on RPA-coated ssDNA.

**Figure 3.**
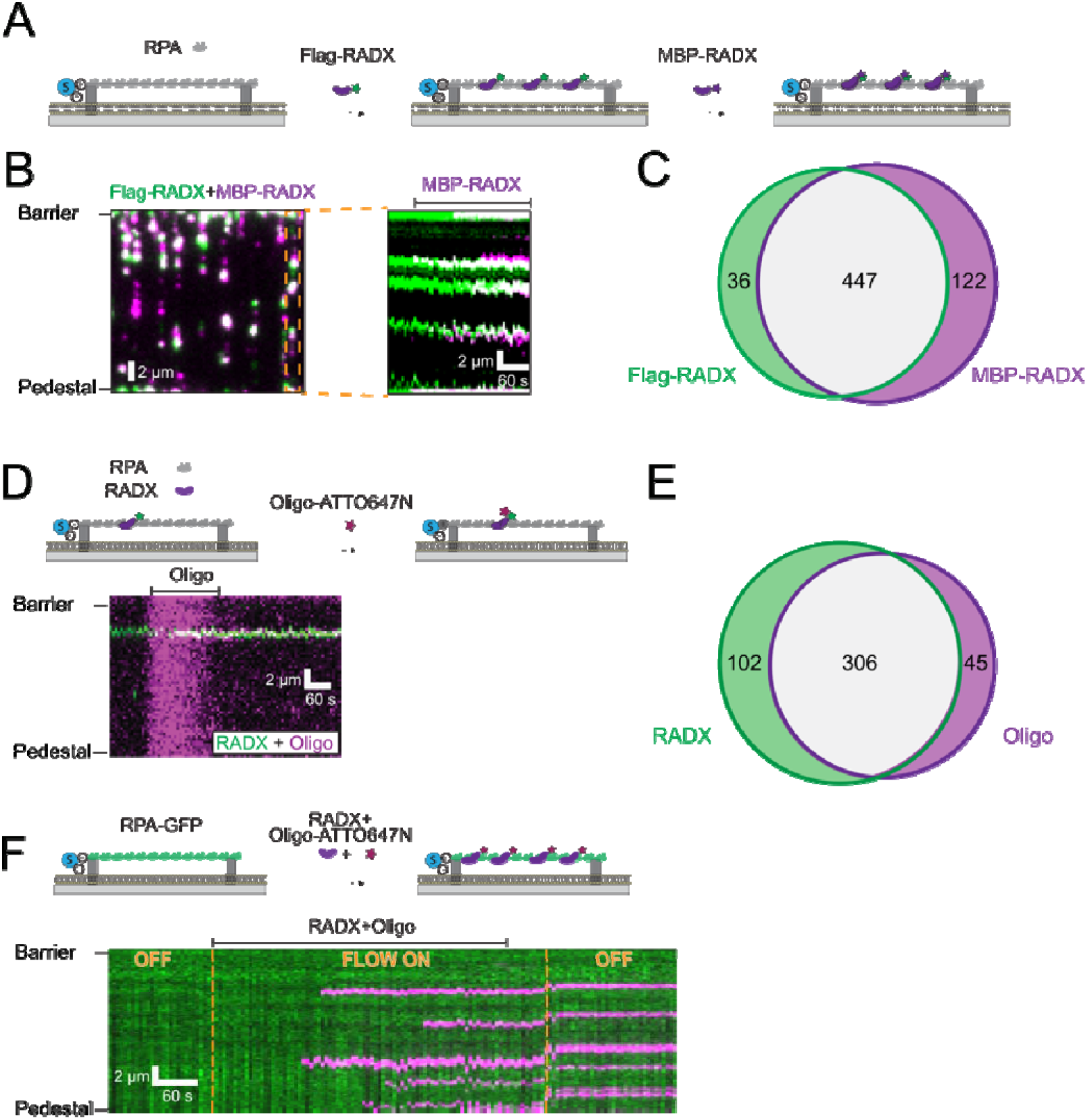
RADX bridges ssDNA via protein-protein interactions. **(A)** Illustration of the double-tethered ssDNA curtain assay. The ssDNA is coated by RPA and anchored between two chromium features above a lipid bilayer. **(B)** Left: RADX forms larger-order assemblies on RPA-coated ssDNA, as imaged via selfassociation of Flag-RADX (green) and MBP-RADX (magenta). Right: Kymograph of one molecule (orange box) indicates that MBP-RADX foci preferentially form at Flag-RADX sites. Flag-RADX was labeled with anti-Flag-Alexa488 and MBP-RADX was visualized with anti-MBP-QD705 antibodies. **(C)** Quantification of MBP-RADX and Flag-RADX co-localization frequency (collected from 113 ssDNA molecules). **(D)** A kymograph indicating that RADX directly captures non-complementary ssDNA oligonucleotides on RPA-coated ssDNA curtains. **(E)** Quantification of RADX and oligo colocalization frequency (collected from 89 ssDNA molecules). **(F)** A kymograph indicating that RADX preincubated with non-complementary ssDNA oligonucleotides efficiently binds on RPA-coated ssDNA curtains. Yellow lines: toggling buffer flow ON and OFF indicates that the oligonucleotide is stably bound on the ssDNA.

Next, we tested whether RADX can capture ssDNA *in trans*. RADX was first incubated with double-tethered ssDNA-RPA curtains, and a fluorescent non-complementary oligo (5 nM) was injected into the flowcell (Figure 3D). Nearly all oligos (87 ± 6%; mean ± SD; N=89 molecules) co-localized with RADX (Figure 3E). The oligos were retained on ssDNA curtains for >10 minutes and were not removed with extensive buffer washes at 100 mM NaCl. Furthermore, pre-incubating RADX with this oligo (2 nM RADX and 1 nM oligo incubated at room temperature for 15 minutes) prior to injection of the mixture in the flowcell also resulted in robust oligo capture on RPA-coated ssDNA curtains (Figure 3F). We did not see any oligos captured on the ssDNA curtains when RADX was omitted from the flowcell (0%; N = 106 ssDNA molecules) (Figure S4E). These data demonstrate that RADX forms multimeric assemblies on RPA-coated substrates. These assemblies can bridge non-complementary ssDNA molecules *in trans* via protein-protein interactions.

### RADX antagonizes RAD51-ssDNA filament formation

RADX is a negative regulator of RAD51 in cells and *in vitro* (16, 17). These cellular results, along with the striking ssDNA compaction observed in our assays, motivated us to examine whether RADX interferes with RAD51 filament assembly. RAD51 nucleation and RAD51-dependent RPA exchange on ssDNA are critical regulatory steps in regulating RAD51 filament formation (33-35). The RAD51-ssDNA filament is over-stretched relative to naked and RPA-coated ssDNA (36). The extent of ssDNA extension serves as a convenient reporter for RAD51 filament assembly and extension (34, 37).

We monitored RAD51 filament formation by measuring the extension of fluorescently end-labeled ssDNA substrates. RAD51 rapidly binds and extends the low complexity ssDNA substrate threefold (Figure 4A, 4B). However, ssDNA that is initially compacted with RADX is no longer extended by RAD51, even when the reaction buffer is supplemented with 2 mM Ca^2+^ to stabilize the RAD51 filament by inhibiting ATPase activity and monomer turnover (38-40) (Figure 4C). Although these data do not rule out that small RAD51 clusters can form on RADX-coated ssDNA, overall the substrate remains compact.

**Figure 4.**
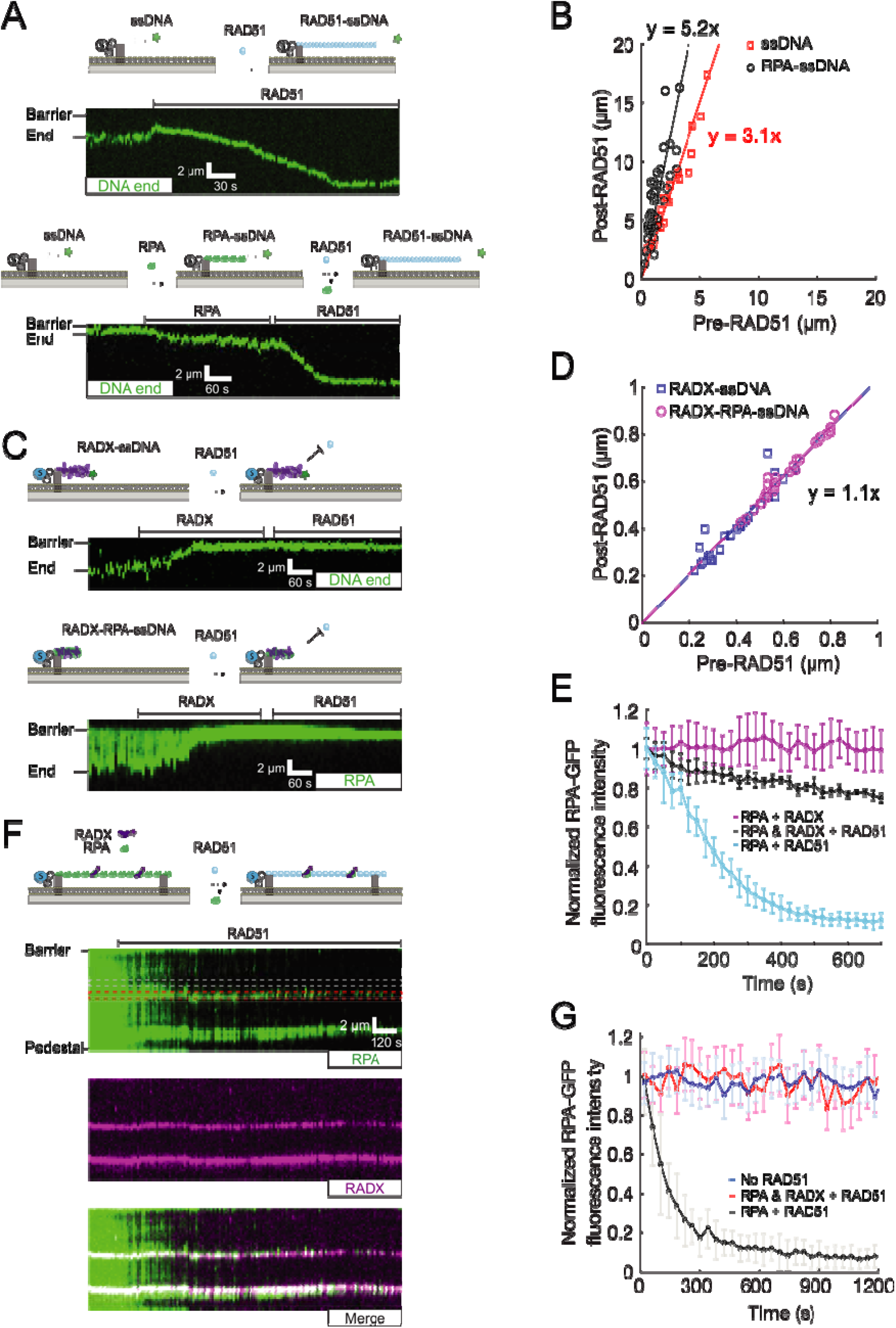
RADX protects RPA from displacement by RAD51 to inhibit RAD51 filament extension. **(A)** Kymographs of RAD51 binding and extending naked (top) or RPA-coated (bottom) ssDNA. **(B)** Quantification of the change in ssDNA length after RAD51 loading (at least 27 ssDNA molecules for the naked and RPA-coated experiments, respectively). **(C)** Kymographs showing that RAD51 is unable to displace RADX and extend ssDNA or RPA-ssDNA. **(D)** Quantification of RADX-compacted SSDNA length after RAD51 is added to the flowcell (at least 21 ssDNA molecules for the naked and RPA-coated experiments, respectively). **(E)** Normalized RPA-GFP fluorescent intensity as a function of time in the presence of RADX (magenta, N=45), RADX and RAD51 (black, N=56), or RAD51 alone (blue, N=40). **(F)** RADX blocks RPA displacement by RAD51 from doubletethered ssDNA. Red box indicates the region where RPA is co-localized with RADX. Grey box indicates an RPA segment without RADX. **(G)** Normalized RPA-GFP fluorescent intensity in the presence of RAD51 (black, N=65), co-localized with RADX in the presence of RAD51 (**red**, N=53), and in the absence of RADX and RAD51 (blue, N=50).

We next tested whether RADX prevents RAD51 loading on RPA-coated filaments. First, we confirmed that 1 μM RAD51 can rapidly replace RPA from ssDNA curtains in imaging buffer containing 2 mM ATP and 2 mM CaCl2 (Figure 4A). As expected, RPA was rapidly replaced by RAD51 along ssDNA, and the ssDNA was extended five-fold (Figure 4B). Injecting 2 nM RADX into the RPA-ssDNA curtains inhibited RAD51 filament formation (Figure 4C). In the presence of RADX, RAD51 cannot extend RPA-ssDNA substrates (Figure 4D), nor can it efficiently displace RPA-GFP from the ssDNA (Figure 4E). To directly observe the dynamics of RPA in the presence of RADX and RAD51, we used double-tethered RPA-coated ssDNA curtains pre-bound with RADX. These curtains were incubated with 1 μM RAD51 in imaging buffer containing 2 mM ATP and 2 mM CaCl2 (Figure 4F). The fluorescence intensity of RPA foci that co-localized with RADX did not decrease, indicating that RADX prevents the removal of RPA by RAD51. In contrast, RPA was rapidly replaced by RAD51 on those segments of the ssDNA substrates that lacked RADX foci (Figure 4G). Taken together, these results demonstrate that RADX inhibits RAD51 filament formation and prevents RPA displacement by RAD51.

## Discussion

Figure 5 summarizes our integrated model for how RADX antagonizes RAD51 activity. RADX uses its three putative OB-folds to bind ssDNA. Protein-protein interactions between RADX monomers assemble the ssDNA substrate into higher-order compacted structures. RADX cannot directly exchange with RPA under the conditions tested in these assays, but sub-saturating RADX binding is sufficient to condense RPA-coated ssDNA and to prevent extensive RAD51 filament assembly. In addition to blocking RPA removal and RAD51 filament assembly, a recent biochemical study also suggested that RADX disassembles pre-formed RAD51 filaments (17). Thus, RADX inhibits RAD51 filament assembly and may also aid in disassembly of preformed RAD51 filaments.

**Figure 5.**
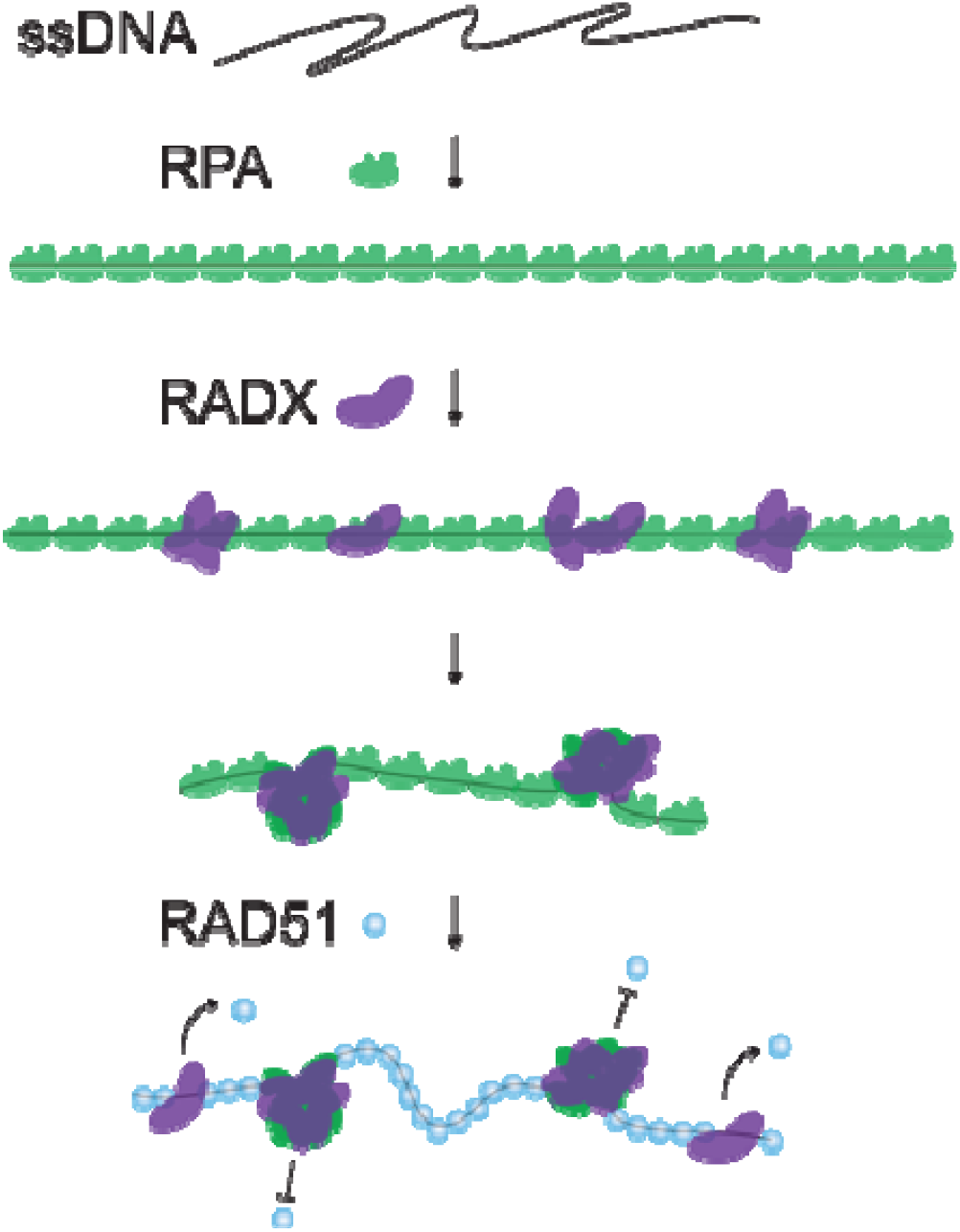
Proposed model of how RADX antagonizes RAD51. RADX compacts RPA-ssDNA filaments, inhibiting RPA displacement and RAD51 filament formation. RADX also removes RAD51 from ssDNA via an unknown mechanism.

Loss of RADX leads to excessive RAD51 activity at stalled replication forks, slowing elongation and causing fork collapse. These studies suggest that RADX antagonizes RAD51 at replication forks to balance fork remodeling and stabilization to maintain genome stability (17). Intriguingly, a recent study also showed that RADX is involved in telomere protection (19). RADX binds exposed single-stranded telomeric DNA along with POT1 to antagonize the accumulation of RAD51 and reduce sister telomere associations. Depletion of either RAD51 or BRCA2 at telomeres rescued RADX depletion, suggesting that RADX also antagonizes homologous recombination in this context (19).

How does RADX stabilize stalled replication forks? Forks that are stalled at lesions are reversed by specialized enzymes to provide time for repair of the lesion (41, 42). However, inappropriate fork reversal can slow fork elongation and result in fork cleavage (43). One possibility is that RADX compacts ssDNA re sulting from dsDNA unwinding during stalled replication, which directly inhibits the formation of RAD51-mediated inappropriate fork reversal. An alternative possibility is that RADX may be involved in fork restoration and may prevent forks from entering the fork protection stage. This stage is characterized by the loading of RAD51 by BRCA2 and the initiation of homologous recombination (15, 44, 45). By blocking RAD51 loading and/or actively dissociating short RAD51 filaments, RADX can antagonize the transition into HR-mediated fork repair. In sum, RADX may be involved in the restoration of fork replication by preventing RAD51 loading and filament formation by condensing ssDNA.

Our observation that RADX forms higher-order oligomers to condense ssDNA raises multiple questions regarding the structural features of this complex and how it is regulated at stalled forks. For example, RADX-ssDNA oligomers need to be disassembled after the lesion is repaired and DNA replication resumes. RADX-ssDNA dissolution can be catalyzed by one or more motor proteins that are required for resuming fork activity (46, 47). For example, BLM helicase may be able to translocate on the ssDNA to strip RADX, akin to its ability to remove RPA and RAD51 from ssDNA (48, 49). Additional possibilities may involve RADX post-translational modifications that either reduce interactions between RADX monomers and/or reduce the affinity of the RADX OB-folds for ssDNA. In direct analogy to RADX, both RAD51 and RPA are phosphorylated and SUMOylated throughout the cell cycle and in response to DNA damage (50–52). Another open question is the interplay between BRCA2/RAD51 and RADX. Cyclin-dependent kinase phosphorylation of the C-terminus of BRCA2 stabilizes RAD51 filaments and regulates fork protection by preventing MRE11-dependent degradation (53, 54). Perhaps BRCA2 can also shift the balance between RADX and RAD51 on ssDNA. Future biophysical and molecular biology studies will need to focus on how RADX forms multi-protein complexes on ssDNA, how these complexes block RAD51, and how these activities are integrated with other fork protection enzymes to restart DNA replication at stalled forks.

## Materials and Methods

### Proteins and Nucleic Acids

Oligonucleotides were purchased from Integrated DNA Technologies (IDT). The plasmids for human wtRPA (pIF47), RPA-GFP (pIF48), and human RAD51 (pIF224) were generous gifts from Dr. Marc Wold and Dr. Mauro Modesti, respectively (20–22). RPA and RAD51 purifications followed previously-published protocols (18, 22, 23).

### RADX purification

All RADX variants were purified from High Five insect cells. The RADX(OB2m) contained the following 10 mutations in the second putative OB-fold: A73S, R240E, R248E, K252E, K255E, K256E, W279A, K304E, R310E, E327A (18). RADX containing an N-terminal Flag tag (pIF434), the RADX(OB2m) mutant (pIF631), and an N-terminal MBP-RADX fusion (pIF632) were expressed by infecting with the appropriate virus for 45 hours following manufacturer-suggested protocols. Pellets were thawed and resuspended in lysis buffer (50 mM Tris-HCl pH 7.5, 500 mM NaCl, 1 mM DTT, and 5% (v/v) glycerol, supplemented with 1× HALT protease cocktail (Thermo Fisher) and 1 mM phenylmethanesulfonyl fluoride (PMSF, Sigma-Aldrich)). After resuspension, cells were homogenized in a Dounce homogenizer (Kimble Chase Kontes) and then centrifuged at 35,000×g for 45 min at 4°C. For Flag-tagged RADX and RADX(OB2m), the supernatant was collected and passed through a column containing 2 mL anti-Flag resin (Sigma-Aldrich F3165) that was pre-equilibrated with lysis buffer. The column was washed extensively with 10 column volumes of wash buffer (20 mM HEPES pH 7.6, 200 mM KCl, 1 mM DTT, 1mM EDTA, 5% (v/v) glycerol) and proteins were eluted with 4 mL of the same buffer but containing 100 μL (5 mg mL^−1^) Flag peptide (Sigma-Aldrich F4799). The eluate was spin concentrated (Sigma-Aldrich CLS431485-251A) and flash-frozen in liquid nitrogen for storage at −80 °C.

For MBP-tagged RADX, the supernatant was collected and passed through a 2 mL Amylose resin (NEB E8021S) pre-equilibrated with lysis buffer. The column was washed with 10 column volumes of wash buffer and proteins were eluted with 8 ml of the same buffer containing 10 mM maltose (Sigma-Aldrich M5895). The eluate was applied to a HiLoad 16/600 Superdex200 pg column (GE Healthcare 28-9893-35) with wash buffer. Peak elution fractions were spin concentrated (Sigma-Aldrich CLS431485-251A) prior to flash freezing in liquid nitrogen and storage at −80 °C. Protein concentration was determined by comparison to a BSA standard curve using SDS-PAGE.

### Preparation of single-stranded DNA substrates

Low-complexity single-stranded DNA substrates were synthesized using rolling circle amplification (24). Briefly, 5 μM of phosphorylated template oligo IF239 and 4.5 μM biotinylated primer oligo IF238 were annealed in T4 ligase reaction buffer (NEB B0202S). The mixture was heated to 75 °C for 5 min and cooled to 4 °C at a rate of −1 °C min^−1^. Annealed circles were ligated with the addition of 1 μL of T4 DNA ligase (NEB M0202S) at room temperature for ~4 hours. Low-complexity ssDNA was synthesized in phi29 DNA polymerase reaction buffer (NEB M0269S), 500 μM dCTP and dTTP (NEB N0446S), 0.2 mg mL^−1^ BSA (NEB B9000S), 10 nM annealed circles, and 100 nM of home-made phi29 DNA polymerase (17). The solution was mixed and immediately injected into the flowcell and incubated at 30°C for ~30 min. ssDNA synthesis was quenched by removing excess nucleotides and polymerase with imaging buffer (100 mM NaCl, 40 mM Tris-HCl pH 8.0, 1 mM MgCl_2_, 1 mM DTT, and 0.2 mg mL^−1^ BSA). All experiments were conducted using the imaging buffer with indicated extra components at 37 °C. When indicated, ssDNA was end-labeled with mouse anti-dsDNA primary antibody (Thermo MA1-35346) followed by an Alexa488-labeled goat anti-mouse secondary antibody (Thermo A28175). For creating double-tethered RPA-coated ssDNA curtains, 2 nM RPA or RPA-GPF in imaging buffer was injected into flowcell at 1 mL min^−1^ flow rate for at least 5 minutes before doing subsequent experiments.

### Single-Molecule Fluorescence Microscopy and Analysis

Flow cells were prepared as previously described (25). Briefly, a 4-mm-wide, 100-μm-high flow channel was constructed between a glass coverslip (VWR 48393 059) and a custom-made flow cell containing 1-2-μm-wide chromium barriers using two-sided tape (3M 665). Single-molecule fluorescent images were collected with a prism TIRF microscopy-based inverted Nikon Ti-E microscope. The sample was illuminated with a 488 nm laser (Coherent Sapphire; 4.1 mW at front prism face) and a 637 nm laser (Coherent OBIS; 20.4 mW at front prism face). Two-color imaging was recorded using dual-electron-multiplying charge-coupled device (EMCCD) cameras (Andor iXon DU897). Subsequent images were exported as uncompressed TIFF stacks for further analysis.

DNA molecules with fluorescently-labeled ends were tracked using a custom-written particle tracking script in FIJI. The resulting trajectories were analyzed in Matlab (Mathworks) to calculate the rate and extent of DNA compaction. For RPA-GFP-coated ssDNA molecules, the GFP intensity was calculated by summing the total pixel intensity over a defined area over every frame using FIJI.

## Acknowledgments

We are grateful to Dr. Miaw-Sheue Tsai and the Expression and Molecular Biology (EMB) Core in Structural Cell Biology of DNA Repair Machines (SBDR) program for providing protein pellets. We thank Dr. David Cortez for expression constructs, preliminary data, and ongoing conversations regarding RADX biology. Dr. Marc Wold and Dr. Mauro Modesti shared RPA and RAD51 over-expression vectors. Finally, we thank members of the Finkelstein lab for carefully reading this manuscript.

## References

1. Cimprich, K.A. and Cortez, D. (2008) ATR: an essential regulator of genome integrity. Nat. Rev. Mol. Cell Biol., 9, 616–627.

2. Byun, T.S., Pacek, M., Yee, M., Walter, J.C. and Cimprich, K.A. (2005) Functional uncoupling of MCM helicase and DNA polymerase activities activates the ATR-dependent checkpoint. Genes Dev., 19, 1040–1052.

3. Myler, L.R., Gallardo, I.F., Soniat, M.M., Deshpande, R.A., Gonzalez, X.B., Kim, Y., Paull, T.T. and Finkelstein, I.J. (2017) Single-Molecule Imaging Reveals How Mre11-Rad50-Nbs1 Initiates DNA Break Repair. Molecular Cell, 67, 891–898.e4.

4. Huertas, P. (2010) DNA resection in eukaryotes: deciding how to fix the break. Nat Struct Mol Biol, 17, 11–16.

5. Bhat, K.P. and Cortez, D. (2018) RPA and RAD51: fork reversal, fork protection, and genome stability. Nature Structural & Molecular Biology, 25, 446–453.

6. Wu, Y., Lu, J. and Kang, T. (2016) Human single-stranded DNA binding proteins: guardians of genome stability. Acta Biochim Biophys Sin (Shanghai), 48, 671–677.

7. Theobald, D.L., Mitton-Fry, R.M. and Wuttke, D.S. (2003) Nucleic Acid Recognition by OB-Fold Proteins. Annual Review of Biophysics and Biomolecular Structure, 32, 115–133.

8. Wold, M.S. (1997) Replication protein A: a heterotrimeric, single-stranded DNA-binding protein required for eukaryotic DNA metabolism. Annu. Rev. Biochem., 66, 61–92.

9. Maréchal, A. and Zou, L. (2015) RPA-coated single-stranded DNA as a platform for post-translational modifications in the DNA damage response. Cell Research, 25, 9–23.

10. Pokhrel, N., Caldwell, C.C., Corless, E.I., Tillison, E.A., Tibbs, J., Jocic, N., Tabei, S.M.A., Wold, M.S., Spies, M. and Antony, E. (2019) Dynamics and selective remodeling of the DNA-binding domains of RPA. Nat. Struct. Mol. Biol., 26, 129–136.

11. Prakash, R., Zhang, Y., Feng, W. and Jasin, M. (2015) Homologous Recombination and Human Health: The Roles of BRCA1, BRCA2, and Associated Proteins. Cold Spring Harb Perspect Biol, 7.

12. Lord, C.J. and Ashworth, A. (2007) RAD51, BRCA2 and DNA repair: a partial resolution. Nature Structural & Molecular Biology, 14, 461 – 462.

13. Davies, A.A., Masson, J.Y., McIlwraith, M.J., Stasiak, A.Z., Stasiak, A., Venkitaraman, A.R. and West, S.C. (2001) Role of BRCA2 in control of the RAD51 recombination and DNA repair protein. Mol. Cell, 7, 273–282.

14. Baumann, P. and West, S.C. (1998) Role of the human RAD51 protein in homologous recombination and double-stranded-break repair. Trends in Biochemical Sciences, 23, 247–251.

15. Zellweger, R., Dalcher, D., Mutreja, K., Berti, M., Schmid, J.A., Herrador, R., Vindigni, A. and Lopes, M. (2015) Rad51-mediated replication fork reversal is a global response to genotoxic treatments in human cells. J Cell Biol, 208, 563–579.

16. Schubert, L., Ho, T., Hoffmann, S., Haahr, P., Guérillon, C. and Mailand, N. (2017) RADX interacts with single-stranded DNA to promote replication fork stability. EMBO reports, 18, 1991–2003.

17. Bhat, K.P., Krishnamoorthy, A., Dungrawala, H., Garcin, E.B., Modesti, M. and Cortez, D. (2018) RADX Modulates RAD51 Activity to Control Replication Fork Protection. Cell Reports, 24, 538–545.

18. Dungrawala, H., Bhat, K.P., Le Meur, R., Chazin, W.J., Ding, X., Sharan, S.K., Wessel, S.R., Sathe, A.A., Zhao, R. and Cortez, D. (2017) RADX Promotes Genome Stability and Modulates Chemosensitivity by Regulating RAD51 at Replication Forks. Molecular Cell, 67, 374–386.e5.

19. Briod, A.-S., Glousker, G. and Lingner, J. (2020) RADX Sustains POT1 Function at Telomeres to Counteract RAD51 Binding, which Triggers Telomere Fragility. bioRxiv, 10.1101/2020.01.20.912634.

20. Henricksen, L.A., Umbricht, C.B. and Wold, M.S. (1994) Recombinant replication protein A: expression, complex formation, and functional characterization. J. Biol. Chem., 269, 11121–11132.

21. Modesti, M., Ristic, D., van der Heijden, T., Dekker, C., van Mameren, J., Peterman, E.J.G., Wuite, G.J.L., Kanaar, R. and Wyman, C. (2007) Fluorescent Human RAD51 Reveals Multiple Nucleation Sites and Filament Segments Tightly Associated along a Single DNA Molecule. Structure, 15, 599–609.

22. Myler, L.R., Gallardo, I.F., Zhou, Y., Gong, F., Yang, S.-H., Wold, M.S., Miller, K.M., Paull, T.T. and Finkelstein, I.J. (2016) Single-molecule imaging reveals the mechanism of Exo1 regulation by singlestranded DNA binding proteins. PNAS, 113, E1170–E1179.

23. Benson, F.E., Stasiak, A. and West, S.C. (1994) Purification and characterization of the human Rad51 protein, an analogue of E. coli RecA. EMBO J, 13, 5764–5771.

24. Schaub, J.M., Zhang, H., Soniat, M.M. and Finkelstein, I.J. (2018) Assessing Protein Dynamics on Low-Complexity Single-Stranded DNA Curtains. Langmuir, 34, 14882–14890.

25. Soniat, M.M., Myler, L.R., Schaub, J.M., Kim, Y., Gallardo, I.F. and Finkelstein, I.J. (2017) Next-Generation DNA Curtains for Single-Molecule Studies of Homologous Recombination. Methods Enzymol, 592, 259–281.

26. Antony, E. and Lohman, T.M. (2019) Dynamics of E. coli single stranded DNA binding (SSB) protein-DNA complexes. Semin. Cell Dev. Biol., 86, 102–111.

27. Hamon, L., Pastré, D., Dupaigne, P., Breton, C.L., Cam, E.L. and Piétrement, O. (2007) High-resolution AFM imaging of singlestranded DNA-binding (SSB) protein—DNA complexes. Nucleic Acids Res, 35, e58.

28. Roy, R., Kozlov, A.G., Lohman, T.M. and Ha, T. (2007) Dynamic Structural Rearrangements Between DNA Binding Modes of E. coli SSB Protein. Journal of Molecular Biology, 369, 1244–1257.

29. Bell, J.C., Liu, B. and Kowalczykowski, S.C. (2015) Imaging and energetics of single SSB-ssDNA molecules reveal intramolecular condensation and insight into RecOR function. eLife, 4, e08646.

30. Chen, R. and Wold, M.S. (2014) Replication Protein A: Single-stranded DNA’s first responder: Dynamic DNA-interactions allow Replication Protein A to direct single-strand DNA intermediates into different pathways for synthesis or repair. Bioessays, 36, 1156–1161.

31. Haring, S.J., Mason, A.C., Binz, S.K. and Wold, M.S. (2008) Cellular Functions of Human RPA1 MULTIPLE ROLES OF DOMAINS IN REPLICATION, REPAIR, AND CHECKPOINTS. J. Biol. Chem., 283, 19095–19111.

32. Gallardo, I.F., Pasupathy, P., Brown, M., Manhart, C.M., Neikirk, D.P., Alani, E. and Finkelstein, I.J. (2015) High-Throughput Universal DNA Curtain Arrays for Single-Molecule Fluorescence Imaging. Langmuir, 31, 10310–10317.

33. Sung, P. and Robberson, D.L. (1995) DNA strand exchange mediated by a RAD51-ssDNA nucleoprotein filament with polarity opposite to that of RecA. Cell, 82, 453–461.

34. Subramanyam, S., Kinz-Thompson, C.D., Gonzalez, R.L. and Spies, M. (2018) Observation and Analysis of RAD51 Nucleation Dynamics at Single-Monomer Resolution. Meth. Enzymol., 600, 201–232.

35. Ma, C.J., Gibb, B., Kwon, Y., Sung, P. and Greene, E.C. (2017) Protein dynamics of human RPA and RAD51 on ssDNA during assembly and disassembly of the RAD51 filament. Nucleic Acids Res, 45, 749–761.

36. Sung, P. and Klein, H. (2006) Mechanism of homologous recombination: mediators and helicases take on regulatory functions. Nature Reviews Molecular Cell Biology, 7, 739–750.

37. Gibb, B., Ye, L.F., Gergoudis, S.C., Kwon, Y., Niu, H., Sung, P. and Greene, E.C. (2014) Concentration-Dependent Exchange of Replication Protein A on Single-Stranded DNA Revealed by Single-Molecule Imaging. PLOS ONE, 9, e87922.

38. Ristic, D., Modesti, M., van der Heijden, T., van Noort, J., Dekker, C., Kanaar, R. and Wyman, C. (2005) Human Rad51 filaments on double- and single-stranded DNA: correlating regular and irregular forms with recombination function. Nucleic Acids Res, 33, 3292–3302.

39. Bugreev, D.V. and Mazin, A.V. (2004) Ca2+ activates human homologous recombination protein Rad51 by modulating its ATPase activity. Proc. Natl. Acad. Sci. U.S.A., 101, 9988–9993.

40. Jayathilaka, K., Sheridan, S.D., Bold, T.D., Bochenska, K., Logan, H.L., Weichselbaum, R.R., Bishop, D.K. and Connell, P.P. (2008) A chemical compound that stimulates the human homologous recombination protein RAD51. Proc. Natl. Acad. Sci. U.S.A., 105, 15848–15853.

41. Atkinson, J. and McGlynn, P. (2009) Replication fork reversal and the maintenance of genome stability. Nucleic Acids Res, 37, 3475–3492.

42. Neelsen, K.J. and Lopes, M. (2015) Replication fork reversal in eukaryotes: from dead end to dynamic response. Nature Reviews Molecular Cell Biology, 16, 207–220.

43. Sogo, J.M., Lopes, M. and Foiani, M. (2002) Fork reversal and ssDNA accumulation at stalled replication forks owing to checkpoint defects. Science, 297, 599–602.

44. Michel, B., Flores, M.-J., Viguera, E., Grompone, G., Seigneur, M. and Bidnenko, V. (2001) Rescue of arrested replication forks by homologous recombination. PNAS, 98, 8181–8188.

45. Subramanyam, S., Jones, W.T., Spies, M. and Spies, M.A. (2013) Contributions of the RAD51 N-terminal domain to BRCA2-RAD51 interaction. Nucleic Acids Res, 41, 9020–9032.

46. Petermann, E. and Helleday, T. (2010) Pathways of mammalian replication fork restart. Nature Reviews Molecular Cell Biology, 11, 683–687.

47. Quinet, A., Lemaçon, D. and Vindigni, A. (2017) Replication Fork Reversal: Players and Guardians. Molecular Cell, 68, 830–833.

48. Patel, D.S., Misenko, S.M., Her, J. and Bunting, S.F. (2017) BLM helicase regulates DNA repair by counteracting RAD51 loading at DNA double-strand break sites. J. Cell Biol., 216, 3521–3534.

49. Yang, J., Bachrati, C.Z., Hickson, I.D. and Brown, G.W. (2012) BLM and RMI1 Alleviate RPA Inhibition of TopoIIIα Decatenase Activity. PLOS ONE, 7, e41208.

50. Dou, H., Huang, C., Singh, M., Carpenter, P.B. and Yeh, E.T.H. (2010) Regulation of DNA Repair through DeSUMOylation and SUMOylation of Replication Protein A Complex. Molecular Cell, 39, 333–345.

51. Shima, H., Suzuki, H., Sun, J., Kono, K., Shi, L., Kinomura, A., Horikoshi, Y., Ikura, T., Ikura, M., Kanaar, R., et al. (2013) Activation of the SUMO modification system is required for the accumulation of RAD51 at sites of DNA damage. J Cell Sci, 126, 5284–5292.

52. Luo, K., Li, L., Li, Y., Wu, C., Yin, Y., Chen, Y., Deng, M., Nowsheen, S., Yuan, J. and Lou, Z. (2016) A phosphorylation–deubiquitination cascade regulates the BRCA2–RAD51 axis in homologous recombination. Genes Dev., 30, 2581–2595.

53. Esashi, F., Christ, N., Gannon, J., Liu, Y., Hunt, T., Jasin, M. and West, S.C. (2005) CDK-dependent phosphorylation of BRCA2 as a regulatory mechanism for recombinational repair. Nature, 434, 598–604.

54. Schlacher, K., Christ, N., Siaud, N., Egashira, A., Wu, H. and Jasin, M. (2011) Double-strand break repair-independent role for BRCA2 in blocking stalled replication fork degradation by MRE11. Cell, 145, 529–542.

