## Supplemental Data for "RADX condenses single-stranded DNA to antagonize RAD51 loading"

SUPPLEMENTARY DATA

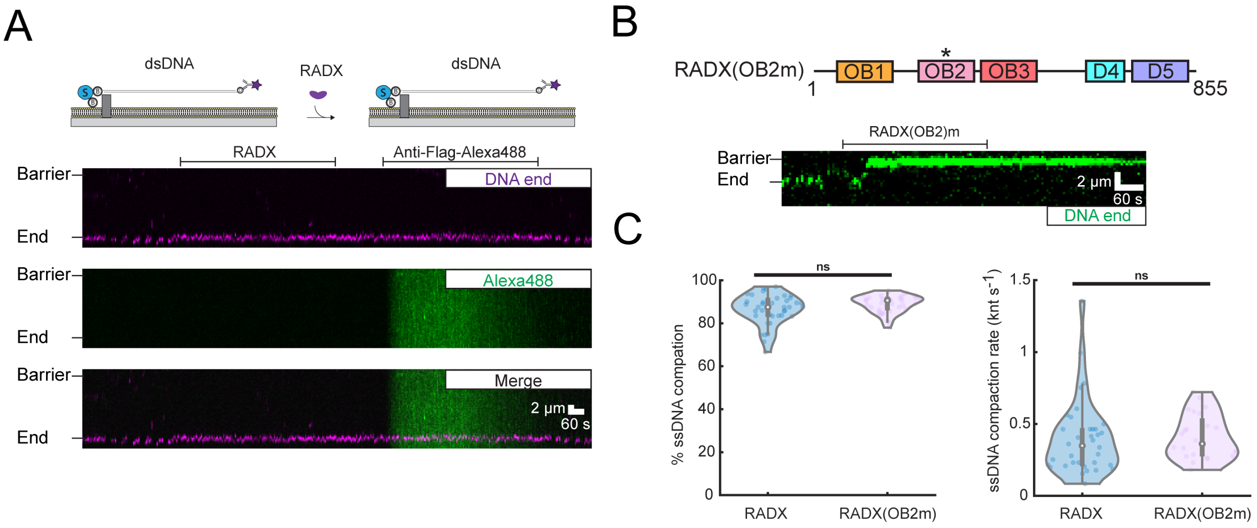


**Figure S1. RADX does not bind to dsDNA. RADX(OB2m) compacts ssDNA.**

(A) A representative kymograph showing RADX does not bind to or compact dsDNA. (B) ssDNA was compacted by RADX(OB2m). (C) ssDNA compaction percentage and rate in the presence of wtRADX and RADX(OB2m).

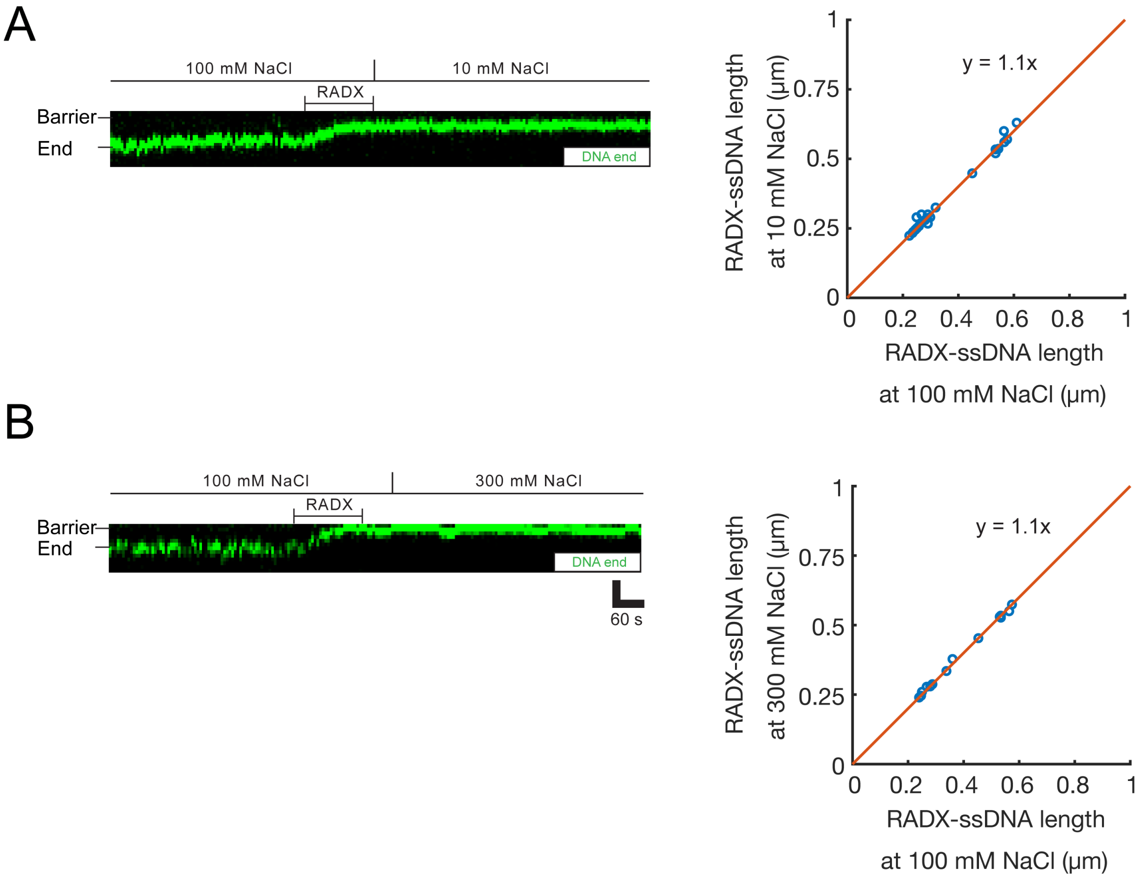


**Figure S2. RADX-ssDNA complexes are insensitive to salt concentrations change.**

(A) A representative kymograph showing NaCl concentration in imaging buffer was switched from 100 mM to 10 mM after ssDNA compacted by RADX. Right panel shows the length of RADX-ssDNA before and after salt concentration switch. (B) A representative kymograph showing NaCl concentration was switched from 100 mM to 300 mM after ssDNA compacted by RADX. Right panel shows the length of RADX-ssDNA before and after salt concentration switch.

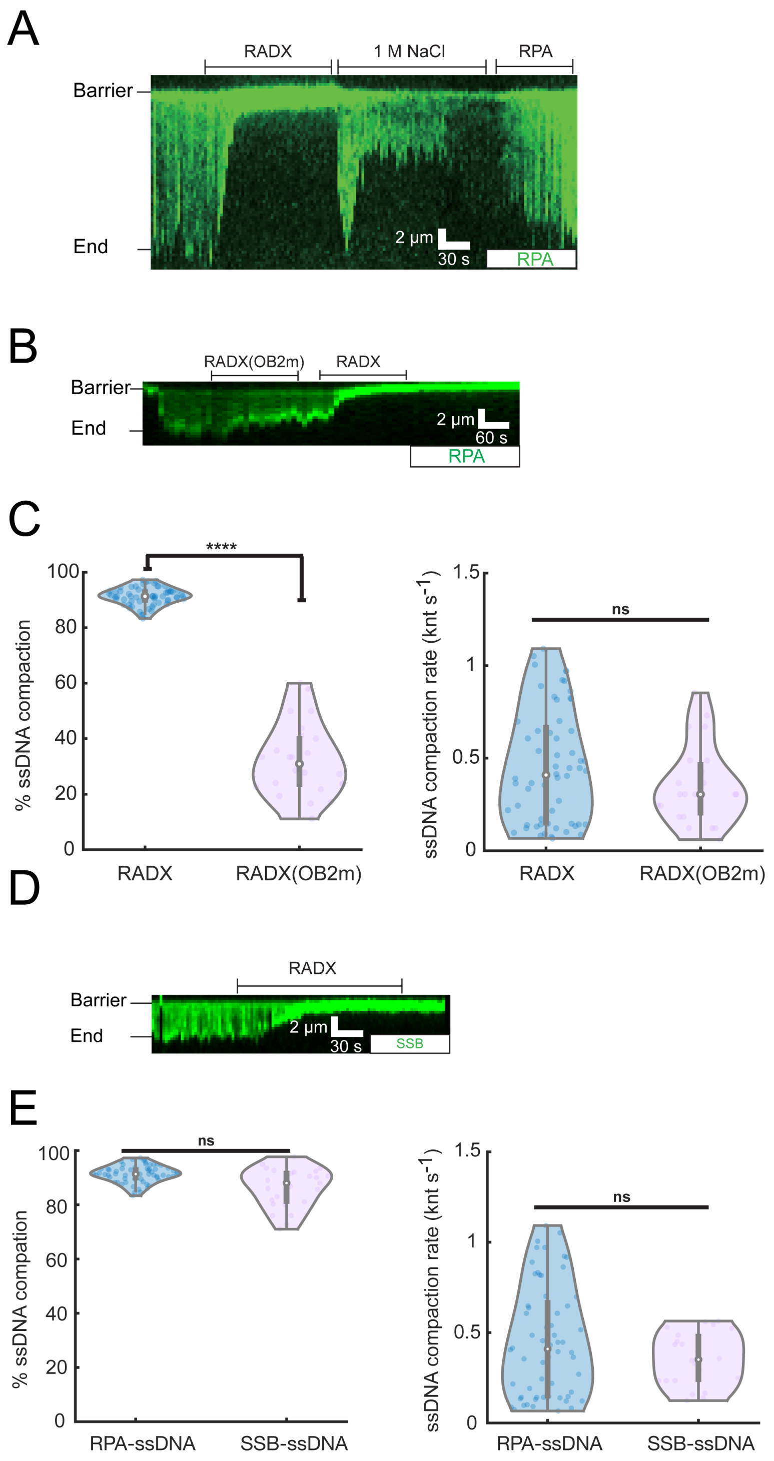


**Figure S3. High salt dissociates RADX from ssDNA. RADX(OB2m) has compromised activity to compact RPA-ssDNA. RADX condenses SSB-ssDNA.**

(A) 1 M NaCl can dissociate RADX from ssDNA and resolve the condensed complexes back to full-length ssDNA molecules. (B) A typical kymograph showing RADX(OB2m) has compromised activity to compact RPA-ssDNA. (C) RPA-ssDNA compaction percentage and rate in the presence of wtRADX and RADX(OB2m). (D) RADX greatly condenses *E.coli* SSB-coated ssDNA. (E) Comparison of SSB-ssDNA and RPA-ssDNA compaction percentage and rate in the presence of RADX.

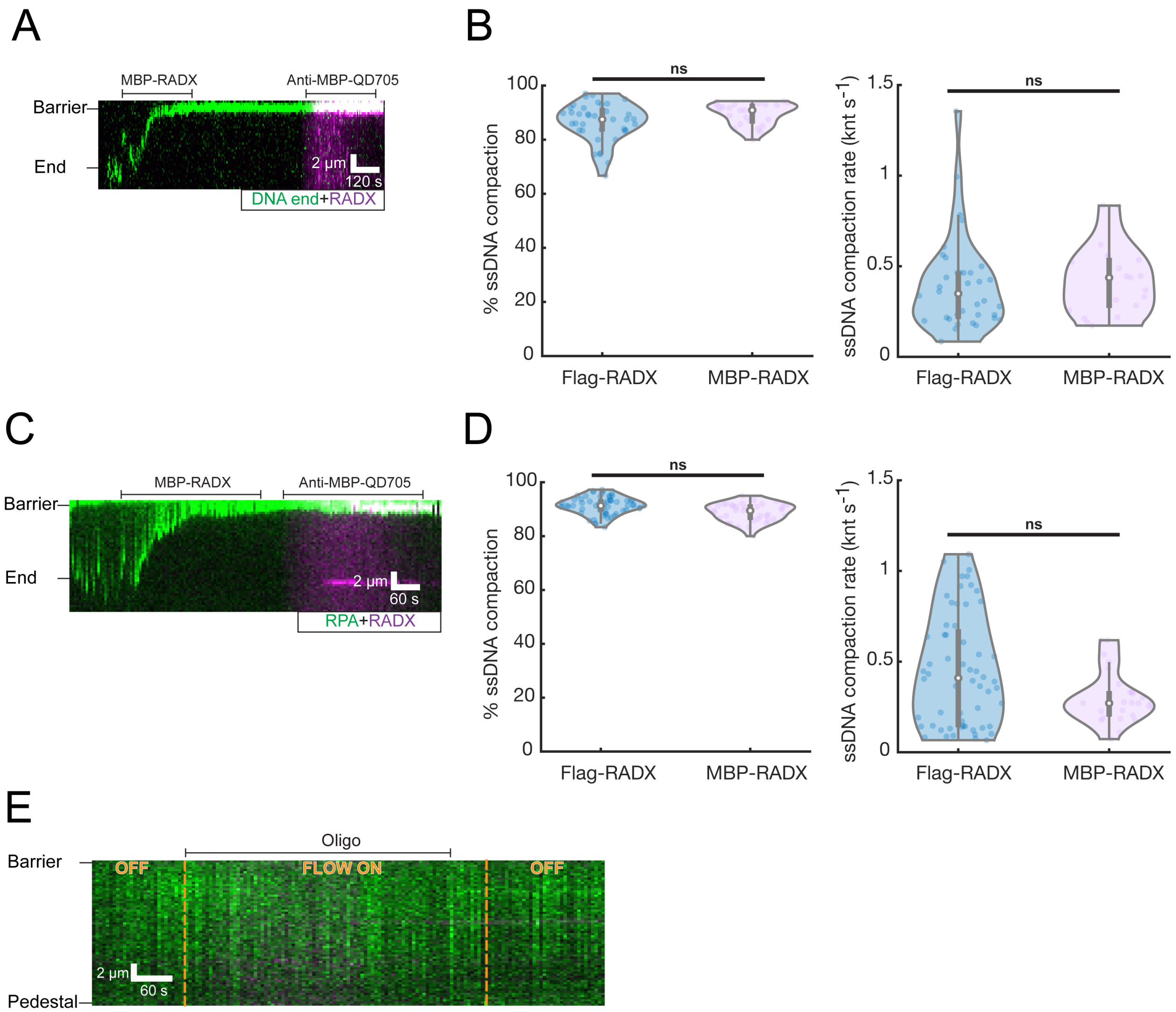


**Figure S4. MBP-RADX is functional comparable with Flag-RADX. Non-complementary oligo does not bind to double-tethered RPA-ssDNA.**

(A) A kymograph showing MBP-RADX condenses ssDNA. (B) ssDNA compaction percentage and rate in the presence of Flag-RADX and MBP-RADX. (C) A kymograph showing MBP-RADX condenses RPA-ssDNA. (D) RPA-ssDNA compaction percentage and rate in the presence of Flag-RADX and MBP-RADX. (E) Non-complementary oligo does not bind to RPA-ssDNA in the absence of RADX.

**TABLE S1. OLIGONUCLEOTIDES USED IN THIS STUDY.**

| Name | Sequence |
|---|---|
| IF327 | GCT GCC GCC CTT GTC ATC |
| IF328 | ACG ATG ACA AGG GCG GCA GCT CCG GAG AGT CTG GGC AAC |
| IF329 | CCT GCA AAG CAC CGG CCT CGT CAG TGG TGG TGG TGG TGG TGA CTA GTA TTT TCA GGA CTG TAA ATC TTG TGA AG |
| IF330 | CAC CAC CAC CAC CAC CAC TGA CGA GGC CGG TGC TTT GCA |
| IF333 | GGA AAA ATC GAA GAA GGT AAA CTG GTA ATC TGG |
| IF334 | CAT ATG TAT ATC TCC TTC TTA AAG TTA AAC AAA ATT ATT TCT AGA GGG G |
| IF238 | 5/Biosg/TC TCC TCC TTC T |
| IF239 | /5Phos/AG GAG AAA AAG AAA AAA AGA AAA GAA GG |
